# Simultaneous Denoising and Baseline Correction of Microplate Raman Spectra Using a Dual-Branch U-Net

**DOI:** 10.64898/2026.04.07.716998

**Authors:** Khaled S. R. Atia, Robert Hunter, Hanan Anis

## Abstract

In this paper, we present a novel dual-branch U-Net architecture for the simultaneous execution of Raman baseline correction and denoising. The network features a shared encoder that diverges into two specialized decoding heads for the Raman signal and for the baseline. The two heads are coupled with a cross-attention gating mechanism. The model offers a way to cross-confirm the peaks by comparing the recovered Raman signal with the baseline corrected spectrum. Moreover, the model offers a new method for quantitative analysis by counting the overall number of photons at a deep Raman decoder block. The model was trained entirely using a custom synthetic data engine explicitly designed to emulate automated HTS acquisitions from microplates via the RamanBot platform. Comprehensive validation demonstrates robust peak recovery on synthetic spectra with signal-to-noise ratios (SNR) as low as 5. Crucially, the model successfully extracts high-fidelity signals from highly noisy glycerol and moderately noisy adenine sulfate experimental samples. Furthermore, quantitative analysis is conducted on guanine samples with different concentrations by counting the Raman photons.

## Introduction

Raman spectroscopy has emerged as a powerful analytical technique across diverse scientific disciplines, including pharmaceutical science,^1^ materials science,^2^ biomedical research,^3^ and environmental science. ^4^ Its widespread adoption is driven by the technique’s ability to provide a label-free, non-destructive molecular fingerprint of a sample. Raman scattering yields highly specific chemical information because it relies on the inelastic scattering of light where the energy shift of scattered photons directly corresponds to the vibrational modes of target molecules.^5^ Despite this analytical power, the throughput of Raman data acquisition continues to lag behind that of other spectroscopic methods. While 96- and 384-well microplates are ubiquitous standards for high-throughput screening (HTS) in fields like genomics and drug discovery, their integration into Raman spectroscopy remains severely restricted. ^6^ A major barrier is the complexity of automated screening. These challenges are typically either mechanical, related to the precision motion systems required for well-by-well screening, or optical, stemming from the difficulties of simultaneous excitation and signal collection. Consequently, commercial high-throughput Raman systems require substantial capital investments, rendering them inaccessible to many research laboratories. To address this challenge, we previously introduced RamanBot,^7^ a cost-effective, automated XYZ Raman acquisition platform. RamanBot utilizes the Cartesian motion system of a commercial 3D printer to sequentially scan samples within a microplate. By replacing the printer’s extruder head with a custom 3D-printed optomechanical mount, the platform effectively houses all the optics necessary to excite and collect Raman signals from individual wells.

While automated platforms like RamanBot resolve the mechanical and optical bottlenecks of high-throughput screening, a fundamental physical barrier remains: the severe signal-to-noise ratio (SNR) challenge inherent to Raman scattering. Only about one in every 10^6^ incident photons scatters inelastically.^8^ Consequently, the inherently weak Raman signal is frequently obscured by two dominant artifacts: high-frequency thermal and shot noise from the detector, and a broad, intense low-frequency baseline primarily caused by sample auto-fluorescence.^9^ This issue is exacerbated in HTS applications, as standard microplates are manufactured from polymers such as polystyrene or polypropylene. These materials generate massive fluorescence signatures that can easily overwhelm the weak signal of the sample within the well. Historically, extracting the true molecular signal has relied on conventional mathematical algorithms. Techniques such as Savitzky-Golay filtering^10^ or continuous wavelet transforms are standard for high-frequency denoising,^11^ while Asymmetric Least Squares (ALS) and Adaptive Iteratively Reweighted Penalized Least Squares (airPLS)^12^ are standard for baseline removal. However, these conventional methods suffer from critical drawbacks. They require fine-tuning of parameters for each sample type, and they routinely fail or distort peak shapes when applied to spectra with extremely low SNRs. Additionally, applying these techniques consecutively leads to loss of peaks or false peaks due to noise. These distortions make tasks such as classification or quantification highly sensitive to the cleanliness of the extracted Raman peaks. To overcome the limitations of classical algorithms, there has been a recent shift toward applying deep learning to Raman spectral processing. Convolutional Neural Networks (CNNs) and Residual Network architectures have been successfully deployed to automate baseline correction and noise reduction.^13–17^ Specifically for denoising, an encoder-decoder network was proposed to enhance the SNR of noisy data,^18^ enabling 10x faster acquisition. Similarly, a generative adversarial CNN was introduced to reduce acquisition time by an order of magnitude for single-cell analysis, ^19^ and a U-Net architecture based on self-supervised learning was constructed for the generic denoising of Raman signals. ^17^ For baseline removal, a residual U-Net^20^ significantly improved the limit of detection (LOD) of Rh6G to 10^−8^ M. Addressing both baseline removal and spectrum denoising simultaneously is often achieved by cascading models, where one model handles the baseline and the other tackles the noise.^14,21,22^ Although successful, this cascaded approach is fundamentally vulnerable to missing subtle peaks or distorting peak intensities. Because the networks operate independently without a shared representational connection, errors generated in the first stage can easily propagate to the second, causing an irreversible loss of chemical information.^23,24^ Moreover, cascaded architectures introduce significant computational and optimization inefficiencies. Deploying two independent models necessitates executing two full forward passes, effectively doubling the inference latency—a severe bottleneck when processing thousands of spectra in high-throughput screening environments.^25^ Furthermore, because the networks are optimized as disjointed entities, they are unable to leverage shared feature representations. ^26^ This isolates the gradient flow, preventing the models from synergistically learning the physical relationship between the low-frequency fluorescence and the sparse Raman scattering, which often leads to suboptimal global convergence. ^27^

To overcome the vulnerabilities inherent in cascaded models, we present a novel two-headed U-Net architecture for simultaneous Raman baseline correction and denoising. This network features a shared encoder, which fundamentally prevents the sequential information loss seen in previous approaches. The encoder is constructed from Runge-Kutta inspired residue blocks^28^ which further reduces the encoder errors. Following the shared bottleneck, the network branches into two distinct decoding heads: one dedicated to extracting the clean Raman signal and the other to recovering the low-frequency baseline. Fundamentally, this dual-head approach operates as a hard-parameter sharing Multi-Task Learning (MTL) framework, enabling the network to leverage shared feature representations rather than relying on isolated gradient flows.^26^ This joint optimization mathematically forces the encoder to learn highly robust, generalized features capable of decoupling the superimposed baseline and Raman signals without destructive interference.

This dual-head approach provides a built-in validation mechanism, allowing the mathematically derived signal (the raw spectrum minus the generated baseline) to be directly compared against the network’s generated Raman signal. Furthermore, the architecture facilitates continuous inter-head communication via a cross-attention gate.^29^ This gate provides spatial “hints” regarding the location of potential peaks by evaluating the differences between the intermediate encoder layers and the baseline’s intermediate layers. By concurrently updating the shared encoder, the two heads work synergistically to ensure no subtle spectral features are missed. Deep supervision is applied to the deep and middle layers of the decoder to ensure the alignment of the features. ^30,31^ The network is trained on a robust synthetic dataset generated by a custom Raman spectrum engine, which was specifically developed to accurately mimic the high-throughput output of RamanBot acquiring samples directly from microplates. To ensure real-world applicability, the model’s performance is subsequently validated on glycerol and adenine sulfate acquired directly from microplates via the RamanBot platform. Finally, we perform quantitative and acquisition time analysis on guanine samples by making use of the Raman photon representation in the Raman branch deep layer.

### Experimental Setup

For data acquisition, we utilized the RamanBot,^7,32^ a previously developed automated screening platform. As illustrated in Fig. 1, the system repurposes the motion gantry of a 3D printer, replacing the standard extrusion head with a custom-designed, 3D-printed optical module. We have previously demonstrated that the RamanBot can autonomously screen microplates by sequentially translating between wells according to a preprogrammed trajectory. This high-throughput capability makes it an ideal platform for rapidly acquiring the large-scale Raman datasets required for machine learning applications.

**Figure 1:**
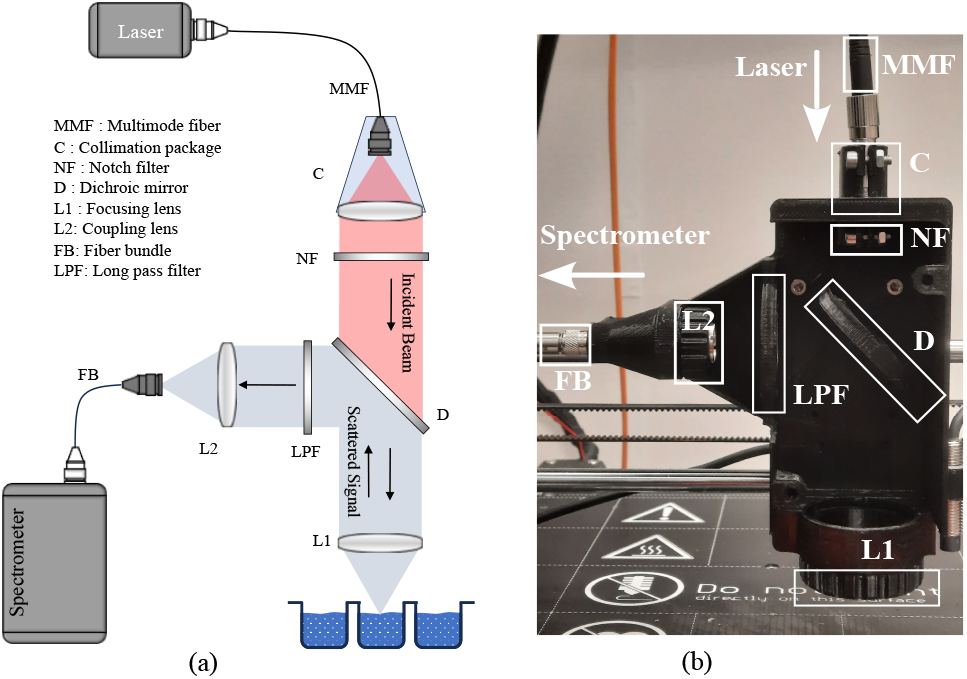
(a) A schematic diagram of the Raman setup. (b) The 3D printed head that mounts all the optics needed to excite wells sequentially.

The internal optical configuration of the head is designed for efficient signal excitation and collection. Excitation light is delivered to the module via a multimode (MM) fiber and collimated by a 0.22 numerical aperture (NA) package. The beam passes through a notch filter to suppress laser sidebands before a 0.4 NA lens focuses the light onto the sample well. The backscattered Raman signal is collected by the same lens and reflected by a dichroic mirror toward the collection path. The signal then passes through a longpass filter to further reject Rayleigh scattering. Finally, a second 0.22 NA lens focuses the filtered Raman signal into a fiber bundle, which routes the collected light to an Andor HoloSpec spectrometer ^33^ for detection.

Figure 2 shows the raw background spectrum captured from one well filled with 330 *µ*L of water. The spectrum was obtained under 1 min acquisition time. This figure shows how complex the background signal of a microplate is. The red dotted line represents the baseline computed using the airPLS technique. ^12^ The inset shows the baseline-corrected Raman spectrum of the microwell. In the first 700 cm^−1^, we noticed an oscillation pattern that can be attributed to the etalon effect. ^34^ This happens due to the multiple reflections between the microwell bottom and the liquid surface.

**Figure 2:**
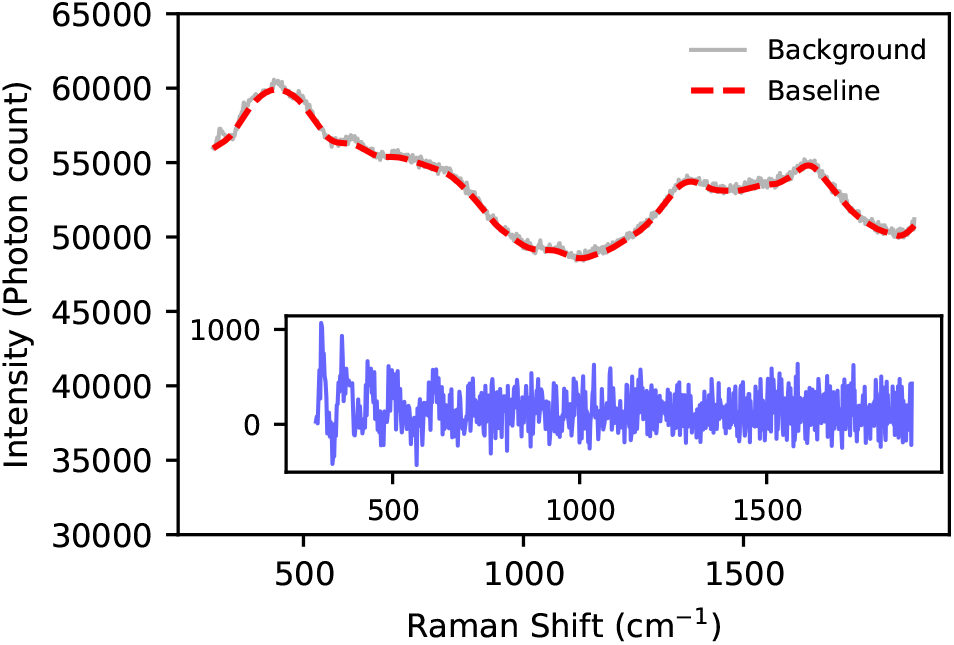
Raw Raman background spectrum and baseline correction obtained from a 96-well plate filled with 330 *µ*L of water. A dashed red line denotes the computed baseline fitted across the raw data. The inset displays the baseline-corrected signal.

## Methodology

### Synthetic Raman Engine

For the deep learning model to achieve optimal performance, it requires training on a substantial volume of high-quality data. Acquiring such a large dataset experimentally is prohibitively time-consuming. Consequently, we developed a synthetic data engine capable of generating extensive spectral datasets that accurately simulate the performance of our experimental setup. The generator models a measured spectrum, *X*(*ν*), as a linear superposition of the Raman signal *S*(*ν*), the fluorescence baseline *B*(*ν*), and the noise *N*(*ν*),^35^ where *ν* represents the Raman shift:

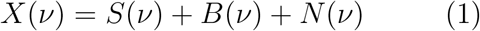

#### Signal Modeling

Because the samples in the microplate are predominantly liquids, their Raman peaks fundamentally exhibit a Lorentzian profile.^8^ However, instrumental broadening introduced by the optical setup and the CCD detector imposes a Gaussian contribution, resulting in faster-decaying tails. ^2^ Therefore, to accurately capture the convolution of natural linewidths and the instrumental response, we model the Raman peaks using a Pseudo-Voigt profile—a which is a linear combination of Gaussian (*G*) and Lorentzian (*L*) functions:

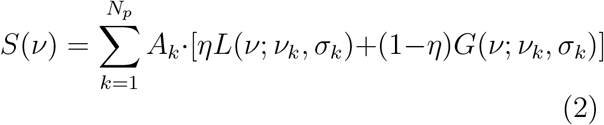

where *N*_*p*_ is the number of peaks, *η* is a weighting parameter that controls the profile shape, *A*_*k*_ is the peak amplitude, and *σ*_*k*_ is the peak width.

#### Baseline

To generate realistic spectral baselines, we empirically collected 1,000 background measurements from a microplate well containing 330 *µ*L of water, utilizing acquisition times ranging from 1 to 60 s. A synthetic baseline is then generated via linear interpolation between two randomly selected experimental background spectra. Furthermore, to simulate phenomena that alter the baseline shape—such as sample evaporation or slight misfocusing—the interpolated baseline is super-imposed with a low-frequency spline curve, *P*_*n*_, constructed from *n* random points distributed across the Raman shift range:

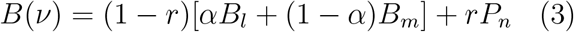

where *α* controls the blending ratio between the two empirical baselines, *B*_*l*_ and *B*_*m*_, while *r* dictates the relative contribution of the synthetic spline to the final baseline.

#### Noise

To accurately model the inherent noise of our experimental setup, we acquired 1,000 dark current spectra using various integration times with the spectrometer shutter closed. Additionally, we collected 1,000 blank spectra across different integration times by focusing the excitation laser 20 cm away from a black microplate to capture ambient and background system noise. To construct a synthetic noise profile, we randomly combine these two empirical noise sources and scale the resulting distribution to achieve the target signal-to-noise ratio (SNR) before superimposing it onto the generated Raman signal.

Figure 3 illustrates three synthetic Raman spectra generated using the proposed pipeline at distinct SNR levels (25, 15, and 5). Each spectrum is synthesized with a randomized combination of peak shifts, amplitudes, and linewidths. Furthermore, the simulated baseline variations across the three generated spectra are clearly observable, reflecting the diverse background conditions accurately captured by our model.

**Figure 3:**
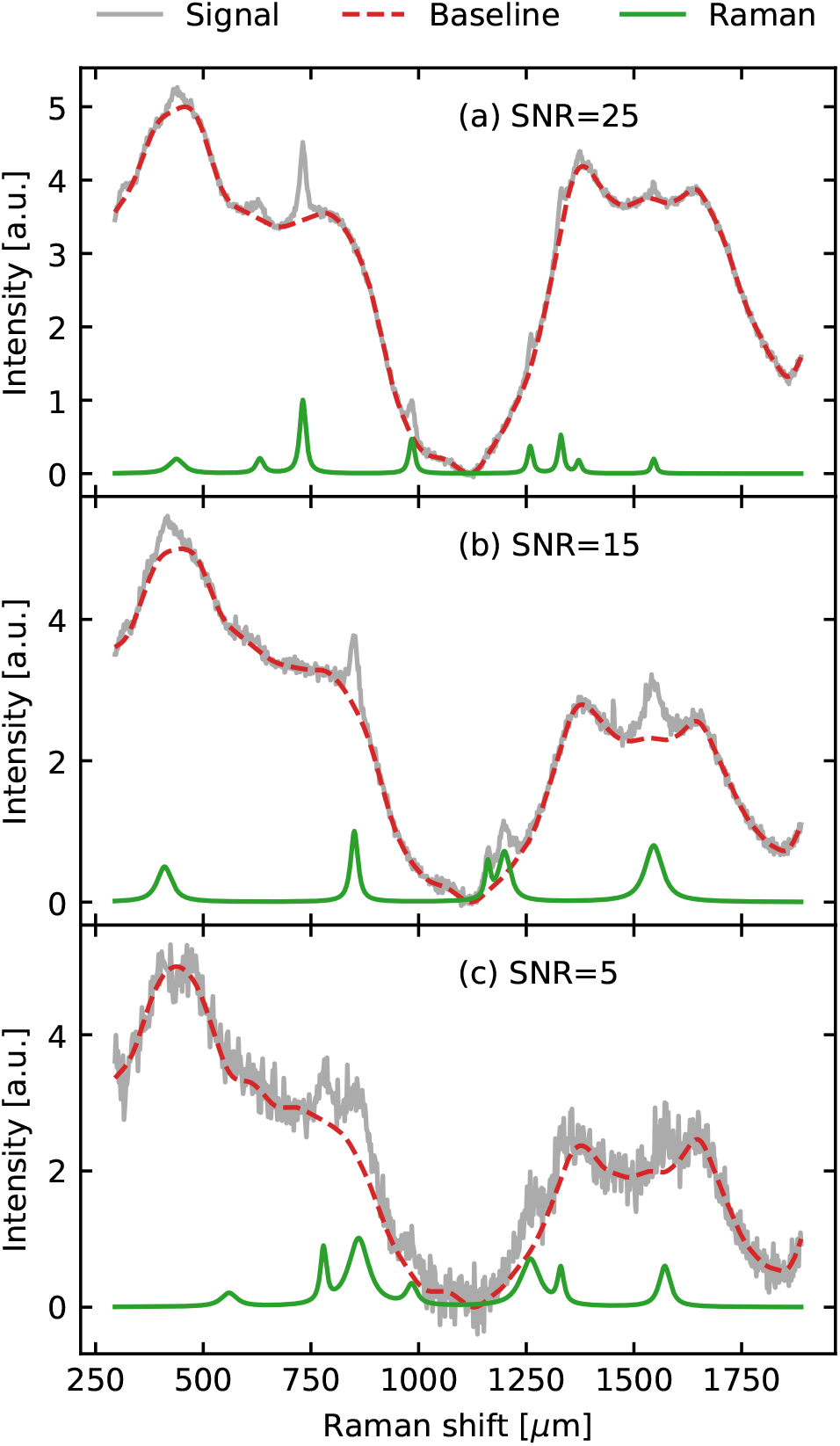
Generation of synthetic Raman spectra under varying noise conditions. The synthetic data were constructed by varying peak parameters (number, width, amplitude) and superimposing a fluorescent baseline. The blue solid line represents the total noisy signal, the red dashed line shows the generated baseline, and the green solid line depicts the pure Raman signal. The panels display representative spectra at Signal-to-Noise Ratios (SNR) of (a) 25, (b) 15, and (c) 5.

#### Training dataset

Table 1 summarizes the parameters used to generate the training datasets. The Raman peak shifts were selected within the range of 300 to 1700 cm^−1^ to prevent spectral truncation at the boundaries of the Raman shift range. The shape parameter, *η*, was chosen to vary between 0.5 and 1.0; this favors a Lorentzian distribution, which is characteristic of the liquid samples used in this study. The spline contribution, *r*, is restricted to the interval [0, 0.4]. This low range reflects the experimental setup involving dark 96-well microplates, where only minimal baseline variations are expected. We used a random number of spline points in the range [3, 10] where we restricted the first and last point to be at the edges of the Raman shift range while the locations of the remaining points were picked randomly. Finally, given the inherently weak nature of Raman signals, we generated a dataset with Signal-to-Noise Ratios (SNR) ranging from 5 to 25. These values were sampled on a logarithmic scale to ensure the model is robustly trained across both low- and high-SNR regimes.

**Table 1:**
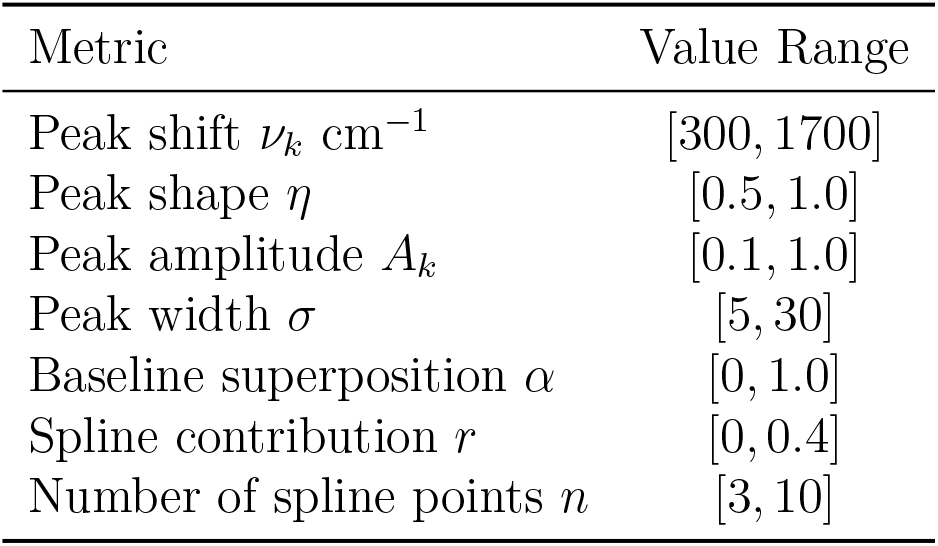
Training dataset parameters.

### Model Architecture

We propose a dual-branch U-Net architecture, illustrated in Fig. 4(a), designed specifically to decouple complex baseline artifacts from weak Raman scattering. The network accepts a 1D Raman spectrum of 864 points following laser line removal. The first residue block expands the single channel to 64 channels. It utilizes a shared encoder composed of three down-sampling blocks that progressively compress the spatial sequence to a length of 108 at the bottleneck while increasing the feature depth to 512 channels. Each down-block is responsible for spatial reduction with advanced ordinary differential equation (ODE) inspired feature extraction.^28^ It consists of a 1D convolution layer with a kernel size of 3 and a stride of 2, which efficiently halves the spatial resolution of the input sequence while maintaining the initial channel depth. The downscaled tensor is immediately normalized using group normalization (hardcoded to 8 groups) to stabilize the learning process without relying on batch statistics, followed by a standard LeakyReLU activation. Finally, the tensor is passed into a Runge-Kutta inspired residue block.^28,36,37^ This expands the feature space to the targeted channels while refining the representations using a 4th-order Runge-Kutta style residual integration, offering enhanced gradient flow and stability compared to standard residue blocks.^37^ Specifically, the block evaluates the tensor through an internal sequence, *F*(*z*), comprising two 1D convolutions (kernel size 3, padding 1) that are each preceded by group normalization and a ReLU activation. The feature representation is then updated using the explicit Runge-Kutta integration step: 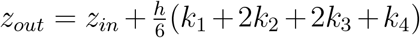, where *h* = 1.0 and the *k* terms represent the four intermediate state evaluations. To account for the low-frequency nature of the expected macroscopic fluorescence, the 512 bottleneck channels are divided asymmetrically: 496 channels for the Raman head and 16 channels for the baseline head. Strict latent space disentanglement between the Raman and baseline channels is enforced through an orthogonal auxiliary loss.^38^ From this latent representation, the architecture branches into two specialized, jointly-trained decoders. The 16 baseline channels and 496 Raman channels are independently up-converted using up-blocks to reconstruct the baseline and Raman signals, respectively. Each up-block is designed to restore spatial resolution while fusing deep semantic features with lower-level spatial details from the encoder. It comprises a learnable up-convolution layer with a kernel size of 2 and a stride of 2, doubling the spatial resolution while halving the channel depth, with padding utilized to guarantee perfect spatial alignment. The corresponding encoder and decoder tensors are concatenated along the channel dimension to bridge their context, then fed into a pre-activated residue block to fuse and refine the features. This residue block consists of two consecutive sequences of group normalization, a ReLU activation, and a 1D convolution (kernel size 3, padding 1). The raw input is added directly to this residual output via an identity skip connection.

**Figure 4:**
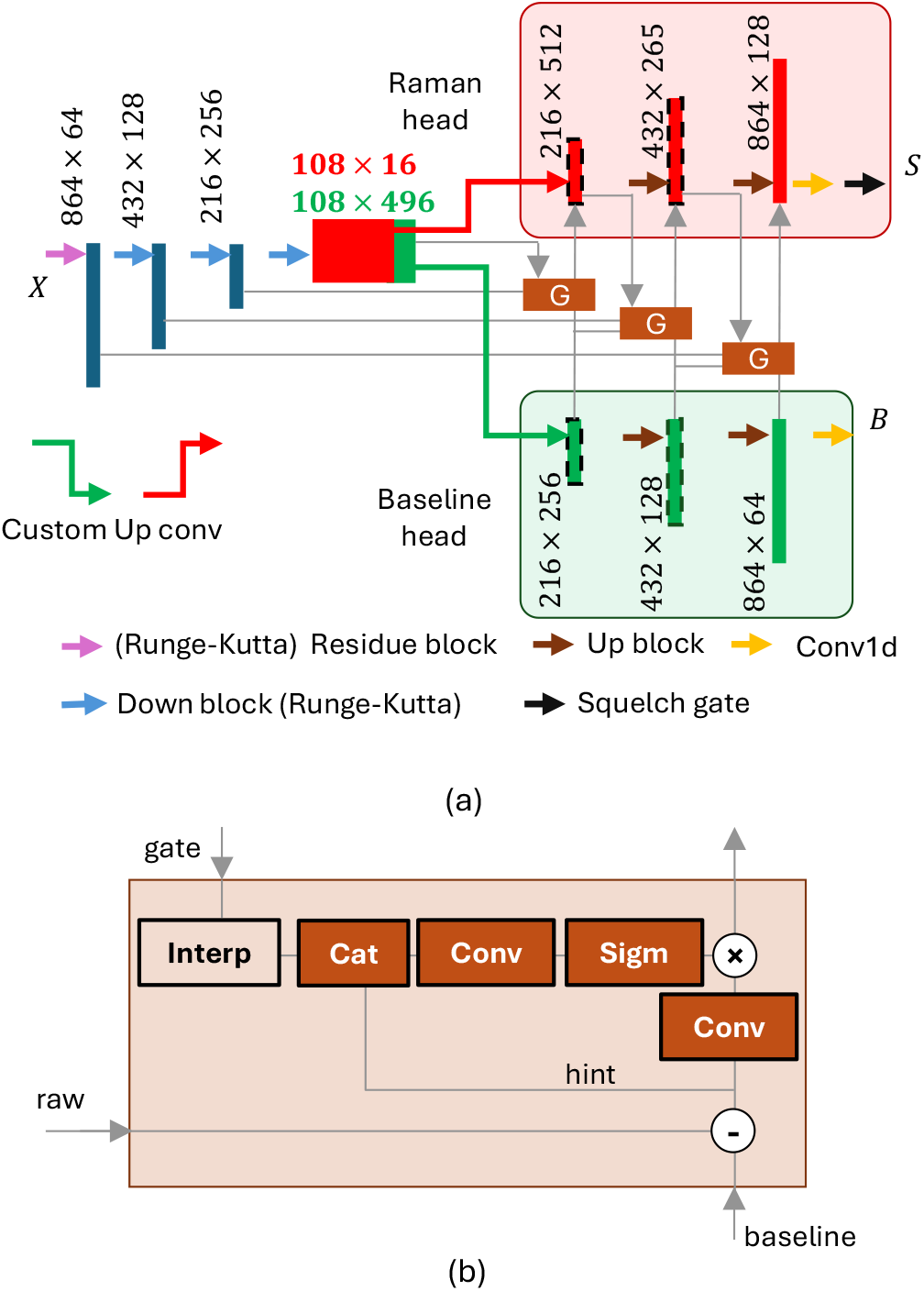
The Dual-Branch U-Net Architecture. The Baseline Head (bottom branch) estimates the background. This estimate is subtracted from the encoder features to create a “Hint” map, which drives the cross attention gate in the Raman Head (top branch), amplifying weak peaks.

The Raman branch communicates with the baseline branch via a cross-attention gate, shown in Fig. 4(b). At each decoder stage, this gate isolates the physical peaks by subtracting the intermediate baseline features from the corresponding high-resolution encoder features. This raw physical “hint” is processed via a 1D convolution to compress the channel dimension. To prevent high-frequency noise from passing through, a deep-semantic gating signal from the previous layer is linearly interpolated and concatenated with the hint signal. A subsequent convolution block and a sigmoid activation function create a binary decision mask used to gate the hint. Consequently, the network enforces a strict cross-checking mechanism where physical evidence is only propagated if corroborated by deep semantic context. To ensure this gating mechanism operates on physically accurate spatial drafts, we incorporate deep supervision^30,31^ at the intermediate decoder layers (highlighted with dotted lines in Fig. 4). These auxiliary losses serve as macroscopic physical anchors. By applying average pooling to downsample the high-resolution ground truth to match the spatial resolution of the intermediate decoders, we enforce strict local energy conservation. This forces the deep, noise-blind layers to correctly map the total photon count and broad geometry of the signals, mathematically constraining the final high-resolution convolutions to only sharpen true physical features rather than memorizing high-frequency shot noise. To ensure the complete suppression of residual noise in regions lacking genuine chemical signatures, a final spatial squelch gate is positioned at the terminal end of the Raman decoder. Constructed from a spatial 1D convolution paired with a sigmoid function, the gate generates a continuous 1D probability mask across the sequence length. By applying this mask to the raw signal prediction, the gate acts as an adaptive filter that squelches inter-peak noise floors to zero while fully opening (1.0) to preserve the mathematically verified Lorentzian geometries of the true Raman peaks. A critical advantage of this dual-branch architecture is the utilization of a shared encoder and a unified bottleneck, fundamentally distinguishing it from conventional cascaded networks. In traditional sequential approaches—where one network models the baseline and a subsequent network performs signal denoising—the system suffers from severe error propagation; if the first network erroneously suppresses a weak Raman peak, the subsequent network cannot recover the lost chemical information. ^39^ By employing a shared encoder, our network constructs a holistic latent representation of the entire optical signal at the bottleneck, preserving the physical context of both the baseline background and the Raman peaks. This ensures the deep Raman layer holds a highly stable, bulk representation of the Raman photons, which is subsequently leveraged for quantitative analysis. Furthermore, branching from a single bottleneck enables joint optimization through multi-task learning. During training, gradients from both the baseline and Raman heads concurrently update the shared encoder, acting as mutual regularizers. This forces the encoder to learn highly robust, generalized feature extractors that perfectly decouple the superimposed signals without destructive interference, while simultaneously reducing the computational overhead compared to deploying independent cascaded networks.

### Loss Function

Our total loss function, ℒ_*total*_, is formulated as a weighted sum of the signal loss (ℒ_*s*_), the baseline loss (ℒ_*b*_), and an orthogonality loss (ℒ_*ortho*_), governed by their respective hyperparameters *w*_*s*_, *w*_*b*_, and *w*_*ortho*_. These loss components are formally defined as follows:

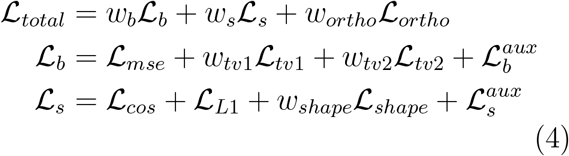

Here, the baseline loss ℒ_*b*_ is composed of a data-fidelity term ℒ_*mse*_—representing the mean squared error (MSE) between the predicted and ground-truth baselines—alongside an auxiliary baseline loss, 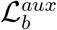. To ensure the predicted baseline reflects the physically slow-varying nature of a typical background, we introduce first- and second-order derivative penalties (ℒ_*tv*1_ and ℒ_*tv*2_), scaled by *w*_*tv*1_ and *w*_*tv*2_, respectively. These regularization terms explicitly penalize abrupt changes in slope and curvature, thereby enforcing spatial smoothness on the generated baseline.

The signal loss, ℒ_*s*_, integrates four components: a cosine similarity error (ℒ_*cos*_), a mean absolute error (MAE) for amplitude fidelity (ℒ_*L*1_), a shape loss (ℒ_*shape*_) representing the MAE between the first derivatives of the predicted and ground-truth Raman signals, and an auxiliary signal loss 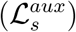. The orthogonality loss, ℒ_*ortho*_, is defined as the squared Frobenius norm of the cross-correlation matrix between the baseline latent channels and the signal latent channels, encouraging optimal feature separation. The auxiliary losses, 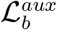 and 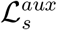, act as intermediate structural anchors that force the network to learn the macroscopic physics of a sample before attempting to resolve high-frequency details. They are expressed as:

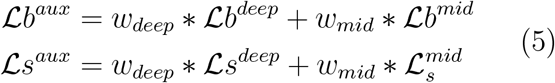

where 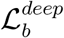 and 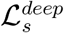 denote the baseline and signal losses at the deep layers, respectively, while 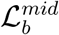 and 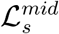 represent the corresponding losses at the intermediate layers. The hyperparameters *w*_*deep*_ and *w*_*mid*_ govern the relative contribution of these deep and middle layers during training. Specifically, the auxiliary baseline loss, 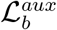, is computed as the mean squared error (MSE) between the target baseline curve and the intermediate predictions of the baseline branch. Conversely, the auxiliary signal loss, 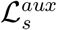, combines the MAE and cosine similarity losses; importantly, it is computed between the ground-truth Raman peaks and the low-resolution spatial maps extracted directly from the deep bottleneck layers of the signal branch.

Table 2 summarizes the hyperparameter values configured for the loss function. The signal and baseline losses are assigned equal weights (*w*_*s*_ and *w*_*b*_), despite the primary focus being the extraction of the inherently weak Raman signal. This balanced weighting is strategic: an accurate baseline prediction—which the network learns readily due to the background’s high amplitude and low-frequency variations—provides a highly reliable hint value for the gating mechanism, ultimately boosting overall model performance. Furthermore, the first- and second-order derivative weights (*w*_*tv*1_ and *w*_*tv*2_) are both set to 0.2 to enforce baseline smoothness while maintaining a tight data fit to the input spectrum. Finally, the shape penalty weight (*w*_*shape*_) is set to 0.5 to rigorously preserve the pseudo-Voigt profile of the detected Raman peaks.

**Table 2:** Loss function parameters.

| Metric | Value |
| --- | --- |
| Signal, baseline weight $w_s, w_b$ | 20,20 |
| Signal shape weight $w_{shape}$ | 0.5 |
| Baseline roughness weight $w_{tv1}$ | 0.2 |
| Baseline curvature weight $w_{tv2}$ | 0.2 |
| Auxiliary deep weight $w_{deep}$ | 1.0 |
| Auxiliary middle weight $w_{mid}$ | 2.0 |
| Bottleneck orthogonality $w_{ortho}$ | 0.5 |

## Training

The network was trained using the Adam optimizer^40^ to minimize the composite multi-task loss function with initial learning rate of 1 × 10^−4^. Training was conducted using a batch size of 64 on a dynamically generated dataset of synthetic spectra. Furthermore, all convolutional weights were initialized using Kaiming normal initialization. Training was conducted on a single NVIDIA RTX 5070 Ti GPU. To ensure stable convergence and prevent gradient explosion across the deep architecture, gradient clipping was enforced with a maximum norm of 1.0. The training process was dynamically modulated using a performance-driven learning rate annealing strategy. Rather than monitoring the total composite loss, the scheduler strictly tracked the validation MAE of the isolated Raman signal prediction, ensuring that learning rate adjustments were driven solely by the network’s quantitative accuracy. If the validation MAE failed to improve for 15 consecutive epochs, the learning rate was reduced by a factor of 0.5 (down to a minimum threshold of 1 × 10^−8^) to facilitate fine weight adjustments near local minima. The validation set comprised 2,000 samples with SNRs logarithmically distributed between 5 and 25. To prevent overfitting and capture the optimal network state, an early stopping mechanism with a patience of 100 epochs was implemented. This mechanism continuously monitored the validation MAE, saving the model weights only when a new global minimum was achieved. The model was trained for a maximum of 1,000 epochs, with early stopping triggered at epoch 759. Upon completion, the training loop automatically restored these best-fit weights, ensuring the final evaluated model represents peak quantitative performance rather than a potentially overfitted state. Figure 5(a) illustrates the progression of the training and validation losses across epochs. The apparent elevation of the training loss relative to the validation loss is a direct result of the assigned hyperparameters and the inclusion of the orthogonality and auxiliary penalties; the validation loss, by contrast, comprises only the unweighted baseline and signal errors. Figure 5(b) details the evolution of the MAE, COS, and shape losses for the Raman branch. The shape loss drops rapidly and smoothly to approximately 0.007, while the COS loss converges more gradually to 0.013, indicating that the network readily learns the overall spectral morphology. Conversely, the MAE decreases slowly to ∼0.02, reflecting the inherent difficulty of accurately reconstructing absolute amplitude values—a challenge stemming from the intrinsically weak nature of Raman peaks relative to the dominant background, particularly at low SNRs. Finally, the orthogonality and auxiliary signal losses are presented in Fig. 5(c). The orthogonality loss drops sharply to ∼0.0003. This minimal value confirms a highly effective disentanglement of the baseline and Raman latent spaces, and its rapid convergence validates that the asymmetrical channel allocation (16 for the baseline head and 496 for the Raman head) provides sufficient representational capacity. Furthermore, the stabilization of the auxiliary signal loss at ∼0.07 suggests that the deep and middle layers of the Raman branch successfully preserve the total photon count and structural integrity of the ground truth, effectively managing the lower spatial resolutions inherent to these intermediate layers.

**Figure 5:**
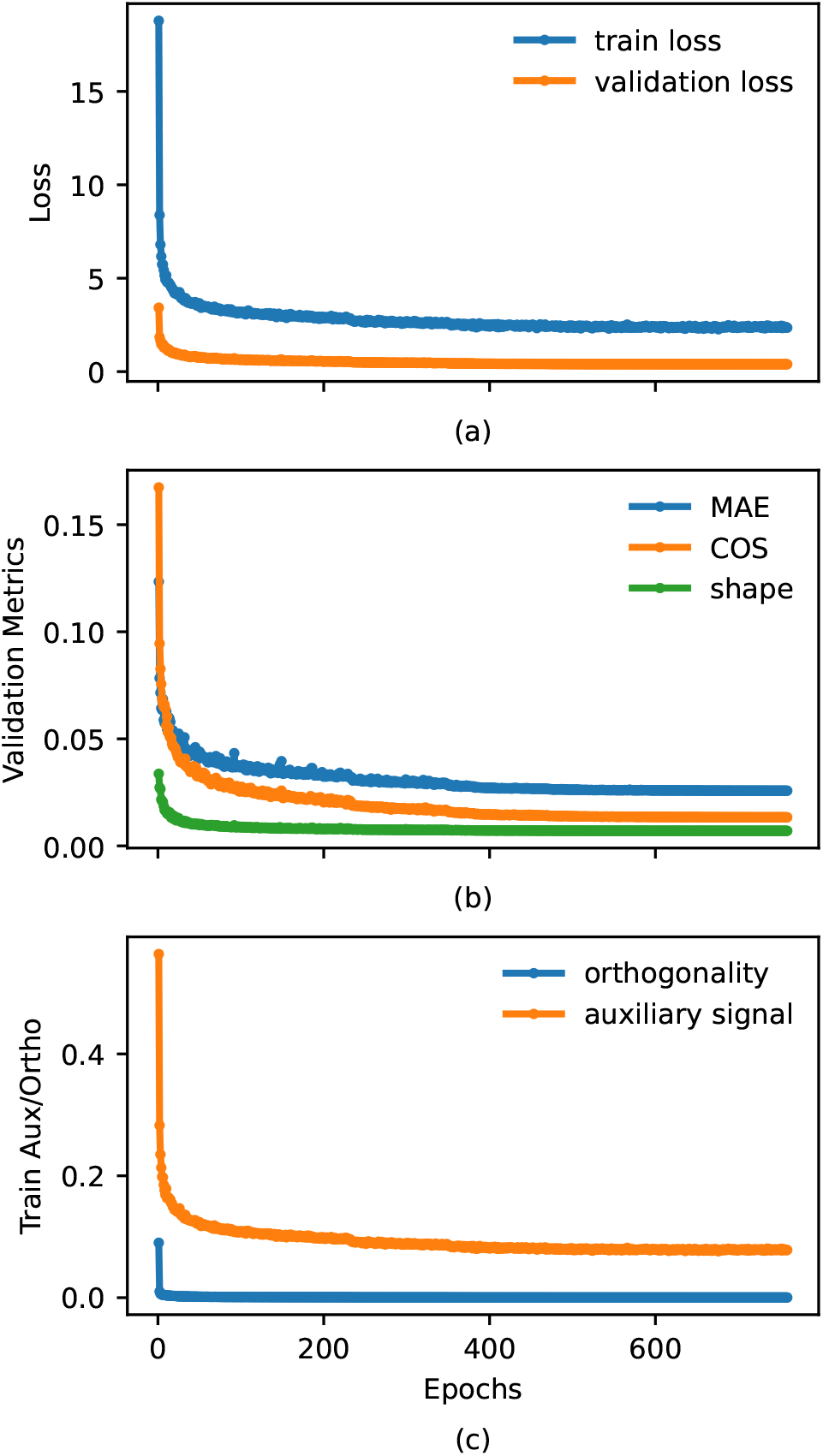
The training loss performance over the epochs broken down into (a) overall training and validation loss, (b) validation metrics (MAE, COS, and shape loss), and (c) Orthogonality and auxiliary loss.

### Validation with Synthetic Data

To systematically quantify the model’s robustness against varying noise levels, we evaluated its predictive accuracy across a continuous range of signal-to-noise ratios (SNRs) from 5 to 25. For this comprehensive analysis, a dedicated test set comprising 500 synthetic spectra was generated for each SNR increment. Figure 6 illustrates the average Mean Squared Error (MSE), Mean Absolute Error (MAE), and Cosine Similarity computed across these datasets. As anticipated, the network’s performance improves monotonically as the SNR increases. The top and middle panels demonstrate that both the MSE and MAE follow similar decay trajectories. At extreme noise levels (SNR = 5), the absolute amplitude errors are at their highest (MSE ≈ 0.005, MAE ≈0.031), reflecting the inherent difficulty of extracting precise intensity values from heavy background interference. However, as the signal quality improves toward SNR 25, these error metrics decrease steadily, reaching minimum values of approximately 0.0015 and 0.017, respectively. This trend highlights the model’s strong quantitative amplitude recovery, particularly as moderate-to-low noise conditions are reached. The bottom panel tracks the Cosine Similarity, which serves as a primary metric for overall spectral shape fidelity. Even under the most severe noise conditions tested (SNR = 5), the model maintains a strong baseline cosine similarity of approximately 0.980, indicating that the core morphological features of the Raman peaks are largely preserved despite the heavy noise. As the SNR increases, this metric climbs rapidly before asymptotically approaching a near-perfect score of ∼0.995 at SNR 25. Collectively, these quantitative trends confirm that the proposed network consistently delivers highly reliable qualitative (shape) and quantitative (amplitude) spectral reconstruction across a broad spectrum of operational conditions.

**Figure 6:**
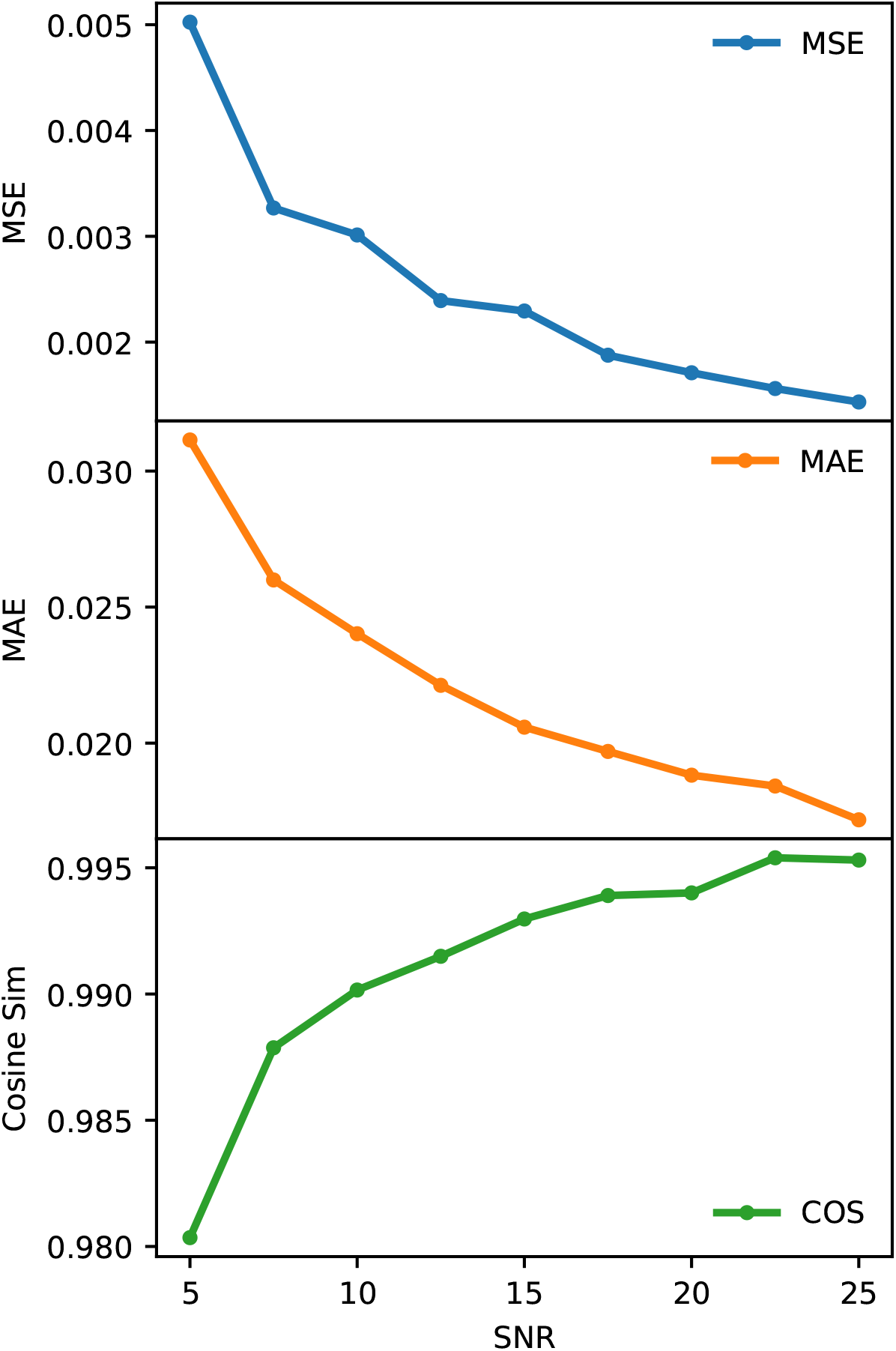
The average MSE, MAE, and cosine similarity computed at different SNR values. A total of 500 spectra were used for each SNR.

To qualitatively evaluate the model’s performance across varying noise levels, we synthesized three representative Raman spectra at signal-to-noise ratios (SNRs) of 5, 15, and 25. Figure 7(a) provides a detailed breakdown of the most challenging case (SNR = 5). This spectrum was constructed with five peaks located at 400, 602, 810, 1300, and 1400 cm^−1^, featuring varying linewidths. Notably, the broad features at 1300 and 1400 cm^−1^ intentionally overlap; this configuration tests the network’s capacity to resolve convoluted peaks and accurately differentiate broad Raman scattering from the underlying background baseline. Additionally, a synthetic cosmic ray spike was injected into a signal-free region at 1680 cm^−1^ to assess the model’s robustness against false-positive detection. The panels in Fig. 7(a) systematically deconstruct the model’s output. The top panel displays the raw synthetic spectrum alongside the model-generated baseline (dotted line), demonstrating a smooth, accurate fit to the slowly varying background trend. The middle panel contrasts the ground-truth Raman signal with the baseline-corrected spectrum. Finally, the bottom panel compares the model’s reconstructed Raman signal directly against the ground truth. The network outputs a recovered spectrum that aligns exceptionally well with the ground-truth signal. Crucially, the model successfully resolves the overlapping peaks, completely rejects the isolated cosmic spike, and exhibits no signs of peak hallucination in the empty regions of the spectrum. Figure 7(b) presents the analysis for the intermediate noise case (SNR = 15). This spectrum incorporates nine distinct peaks—located at 380, 405, 560, 612, 812, 905, 1050, 1300, and 1720 cm^−1^—which exhibit varying linewidths and multiple overlapping regions. As shown in the top panel, the network-generated baseline smoothly tracks the background trend, passing cleanly beneath the prominent Raman features without being skewed by their high intensities. The middle panel compares the baseline-corrected spectrum with the ground truth, illustrating that the relative peak strengths are robustly maintained despite the background removal. Finally, the bottom panel reveals excellent agreement between the model’s reconstructed Raman signal and the ground-truth data. The network accurately preserves both the amplitude heights and the peak widths, confirming its high fidelity in spectral recovery at moderate noise levels. Finally, the low-noise case (SNR = 25) is demonstrated in Fig. 7(c). This spectrum is constructed with 12 Raman peaks located at 360, 405, 560, 612, 735, 805, 901, 1070, 1290, 1480, 1610, and 1730 cm^−1^, featuring varying linewidths and multiple overlaps. To test the model’s ability to recover signals in extreme edge cases, a synthetic hot pixel was intentionally inserted so that it directly overlaps with the 1480 cm^−1^ peak. Consistent with the previous cases, the top panel shows that the generated baseline perfectly tracks the slowly varying behavior of the raw spectrum, remaining unaffected by the strong peaks and heavily overlapped regions. The middle panel contrasts the baseline-corrected spectrum with the ground truth, where a precise match in peak height and width can be observed. The bottom panel demonstrates an excellent match between the reconstructed Raman spectrum and the ground truth. Notably, the generated Raman signal efficiently filters out the hot pixel present at the 1480 cm^−1^ peak while preserving the underlying true Raman feature.

**Figure 7:**
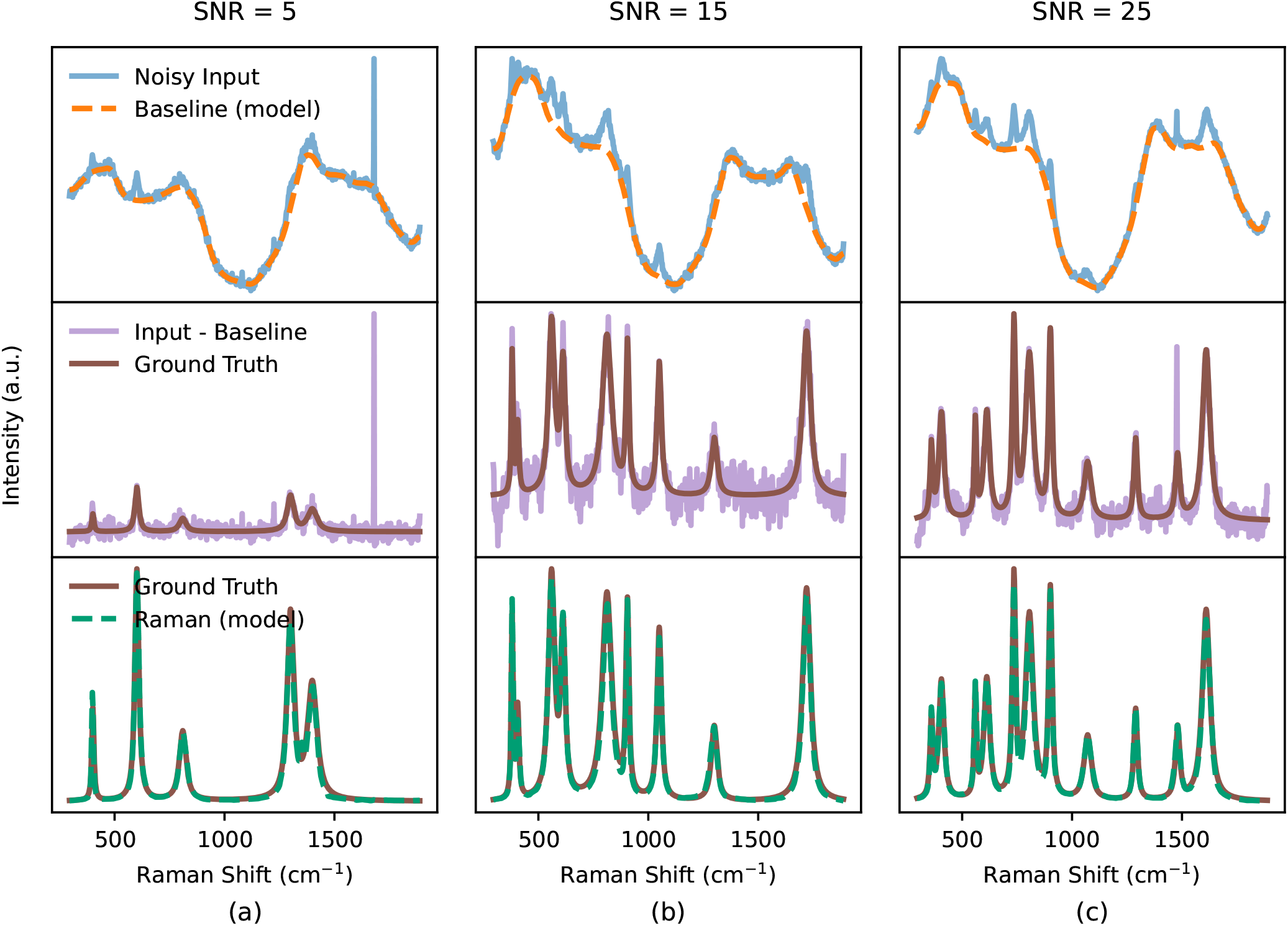
Denoising performance on synthetic data. Top row: Input noisy spectra SNR of 5, 15, and 25. Middle row: Baseline corrected spectrum using the baseline generated by the model overlaid on the ground truth. Bottom row: Comparison between the ground truth and the recovered Raman signal.

Complementing the qualitative visual inspection, Table 3 summarizes the model’s quantitative performance across the three simulated noise levels using Mean Squared Error (MSE) and cosine similarity metrics. The cosine similarity scores demonstrate an exceptionally high degree of spectral fidelity, maintaining a value of 0.996 even under the most severe noise conditions (SNR = 5). As expected, this metric steadily improves as noise decreases, reaching 0.9981 at SNR 25. This trend quantitatively confirms the network’s robust capability to preserve the complex morphological structure and relative peak ratios of the underlying Raman signatures. Concurrently, the MSE values remain consistently constrained to the order of 10^−4^ across all tested noise tiers. These remarkably low error magnitudes indicate that the model not only recovers the correct spectral shape but also accurately reconstructs the absolute intensity values of the Raman signal, maintaining strong amplitude fidelity regardless of the background noise severity.

**Table 3:** MSE and cosine similarity.

| SNR | MSE | Cosine similarity |
| --- | --- | --- |
| 5 | $2.7 \times 10^{-4}$ | 0.996 |
| 15 | $5 \times 10^{-4}$ | 0.9976 |
| 25 | $3.6 \times 10^{-4}$ | 0.9981 |

### Validation with Real Data

To evaluate the model’s generalization capabilities, it was applied to real experimental Raman spectra. A critical aspect of these results is the demonstration of a highly successful ‘sim-to-real’ transfer. It must be emphasized that the network was trained exclusively on the synthetic data engine described previously, without any exposure to real experimental noise profiles, detector artifacts, or genuine auto-fluorescence. Because the dual-branch architecture inherently respects the physical constraints of Raman scattering, it successfully generalizes to the complex optical realities of true experimental samples. This capability effectively eliminates the bottleneck of acquiring massive, labeled experimental datasets and bypasses the need for sample-specific transfer learning. The model was initially tested on aqueous glycerol solutions at concentrations of 100, 150, and 200 mM. For each measurement, a 330 *µ*L aliquot was dispensed into a 96-well plate, and the laser focus was optimized at the sample surface. Figure 8(a) presents the raw acquired spectrum, which is heavily dominated by the background baseline, obscuring most Raman features. Application of the proposed model yielded the baseline fit (dotted line) shown in Fig. 8(a). Although the network was not explicitly exposed to this complex background during training, it successfully generates a smooth fit that accurately avoids the Raman peaks. Figure 8(b) compares the Raman signal recovered by our model’s Raman head with conventional processing methods—specifically, baseline removal via airPLS^12^ followed by Savitzky-Golay smoothing. ^10^ The results shown are for the 100 mM concentration. The airPLS-treated spectrum retains excessive noise, hindering peak identification, and subsequent application of the smoothing filter fails to sufficiently clarify the true Raman features. In contrast, the signal processed by our model exhibits minimal noise, yielding clearly distinguishable peaks.

**Figure 8:**
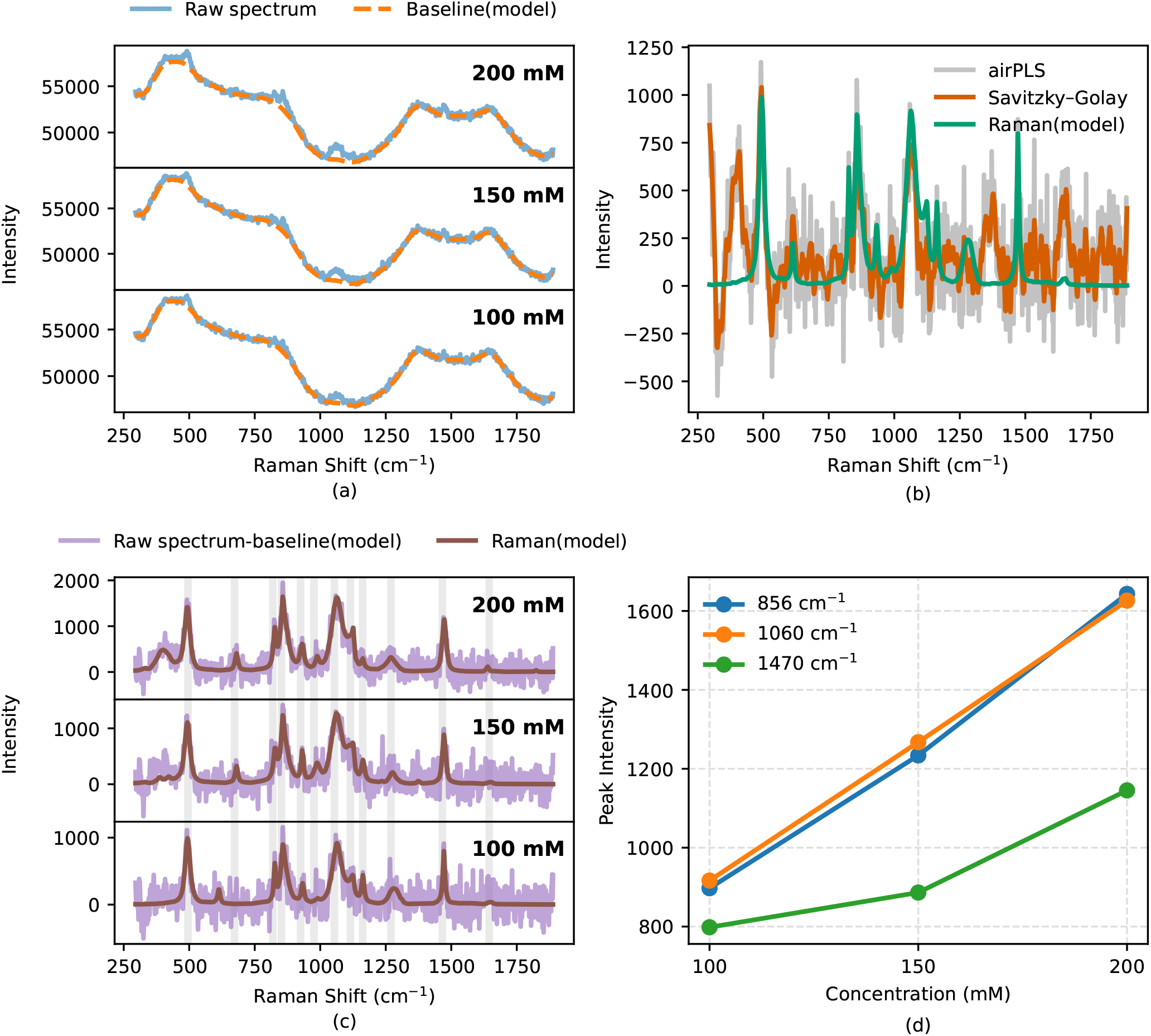
Denoising performance on glycerol samples. (a) Acquired raw spectra for different concentrations 100, 150, 200 mM compared against the recovered baseline. (b) Comparison between the model Raman signal, the baseline corrected spectrum using airPLS, and the smoothed Raman signal using Savitzky-Golay filter. (c) The recovered Raman signal in comparison with baseline corrected spectrum using our model. The shaded columns represent the reported peaks at 495, 673, 819, 852, 925, 976, 1054, 1115, 1162, 1270, 1466, and 1645 cm^−1^. (d) The three dominant peaks tracked for the different concentrations.

Figure 8(c) contrasts the difference signal (raw spectrum minus generated baseline) with the model-recovered Raman signal across the three concentrations. This difference signal represents the raw Raman signature buried under substantial noise. The network-recovered signal demonstrates distinct peaks that align accurately with the underlying features in the difference signal. Key detected peaks are located at 495, 673, 819, 852, 925, 976, 1054, 1115, 1162, 1270, 1466, and 1645 cm^−1^. The peaks at 976 and 673 cm^−1^ correspond to CH_2_ rocking modes and C-C bending vibrations, respectively. Additionally, the broad bands from 800–900 cm^−1^ and 1000–1150 cm^−1^ are attributed to in-phase and out-of-phase alcohol group vibrations, respectively. ^41^ The feature at 1645 cm^−1^ arises from the H-O-H bending mode of the aqueous solvent. The remaining spectral features are consistent with previously reported assignments for glycerol.^41–43^ Finally, the intensities of three prominent peaks (856, 1060, and 1470 cm^−1^) were tracked across the tested concentrations (Fig. 8(d)). Peak intensities correlate positively with concentration; notably, the 856 and 1060 cm^−1^ peaks exhibit highly linear responses suitable for quantitative analysis, whereas the 1470 cm^−1^ peak demonstrates a slight deviation from linearity.

To further evaluate the model’s performance, it was applied to moderately noisy spectra. For this analysis, three different concentrations of adenine sulfate (30, 40, and 50 mM) were prepared in 1 M sodium hydroxide. Figure 9(a) presents the collected raw spectra for these concentrations alongside the baseline generated by our proposed model. The model accurately tracks baseline variations, particularly beneath prominent Raman peaks. Figure 9(b) compares the Raman signal recovered by our model with conventional airPLS baseline correction and subsequent Savitzky-Golay smoothing for the 30 mM sample. Notably, the airPLS-corrected spectrum exhibits negative intensities, indicating an overfitted baseline. This overfitting propagates to the smoothed signal, compromising the underlying data despite the filter’s denoising effects. In contrast, Figure 9(c) demonstrates that the model-generated Raman signal closely aligns with the difference signal (raw spectrum minus generated baseline). Furthermore, key Raman peaks were successfully detected at 545, 633, 730, 988, 1075, 1135, 1205, 1257, 1334, 1372, 1450, and 1548 cm^−1^. The dominant peak at 730 cm^−1^ corresponds to the highly polarized, characteristic ring-breathing mode of the adenine ring.^44^ Additionally, the strong peak at 1334 cm^−1^ is attributed to the external C-NH_2_ stretching vibration, while the symmetric stretching of the sulfate ion is evidenced by the peak at 988 cm^−1^.^45^ The remaining features are consistent with previously reported assignments.^44,46^ Finally, tracking the intensities of the 730, 988, and 1334 cm^−1^ peaks across the three concentrations reveals a robust linear trend, confirming the model’s reliability for quantitative analysis. Overall, these results demonstrate a clean extraction of the sparse Raman signature from the broad baseline.

**Figure 9:**
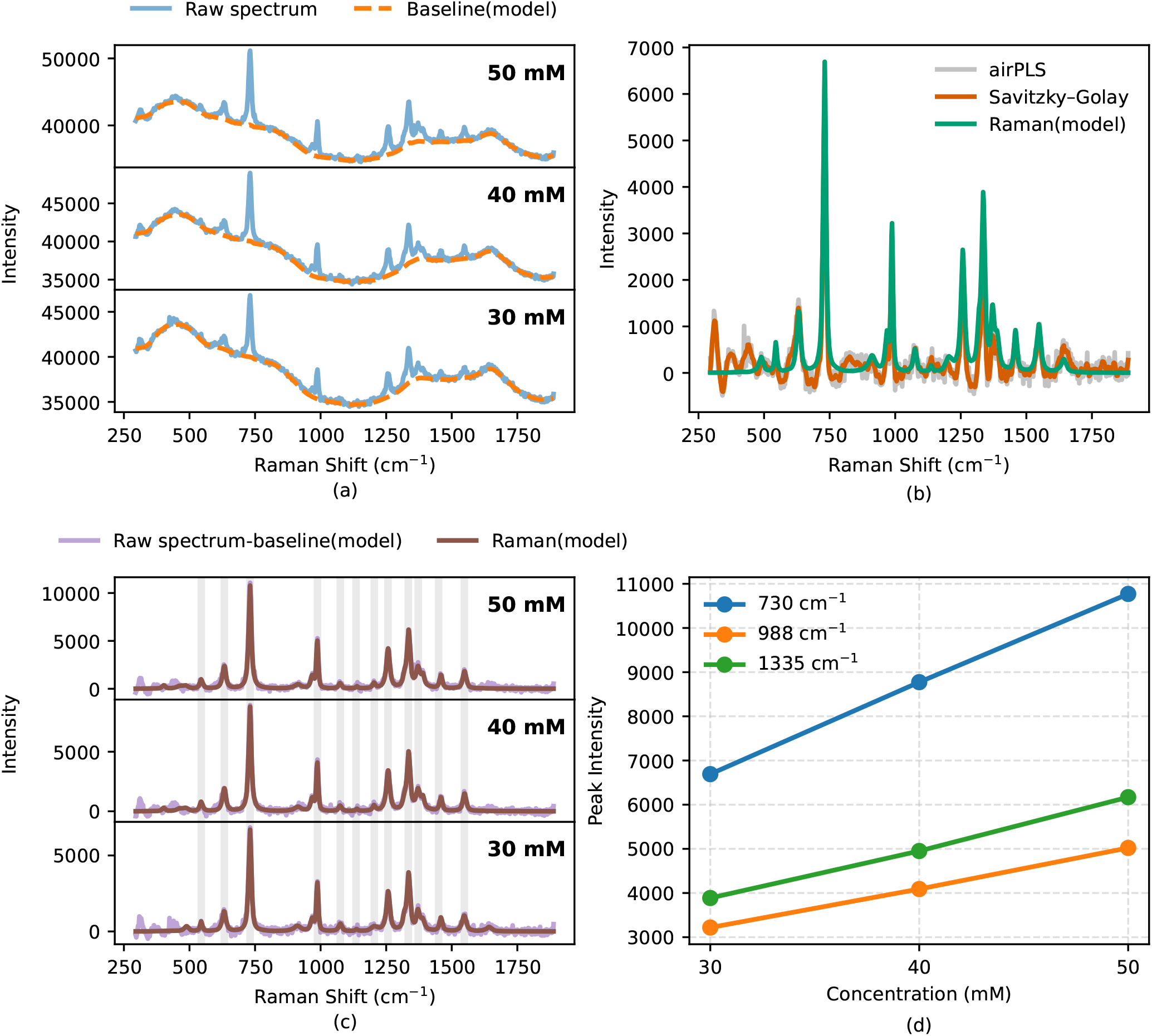
Denoising performance on adenine sulfate samples. (a) Acquired raw spectrum for different concentrations 30, 40, 50 mM compared against the recovered baseline. (b) Comparison between the model Raman signal, the baseline corrected spectrum using airPLS, and the smoothed Raman signal using Savitzky-Golay filter. (c) The recovered Raman signal in comparison with baseline corrected spectrum using our model. The shaded columns represent the reported peaks at 545, 633, 731, 988, 1075, 1135, 1205, 1257, 1334, 1372, 1450, and 1548 cm^−1^. (d) The three major peaks tracked for the different concentrations.

To further investigate the model’s performance in quantitative analysis, we prepared seven different concentrations (ranging from 20 to 80 mM) of guanine diluted in 1 M NaOH. Figure 10(a) illustrates the model’s performance on a subset of these concentrations (30, 40, and 50 mM). The validity of the peaks detected by the Raman head is confirmed by their alignment with the difference signal (raw spectrum minus recovered baseline). The model successfully recovered major peaks at 654, 1165, 1200, 1230, 1270, 1335, 1380, 1460, and 1540 cm^−1^ (highlighted in Fig. 10(a)). The peak at 654 cm^−1^ represents the purine ring breathing mode. Additionally, the peak at 1230 cm^−1^ corresponds to the C2-NH_2_ stretching mode, while the feature at 1380 cm^−1^ represents the C2-N3 stretching mode. The remaining peaks are consistent with previously reported assignments.^47,48^ Monitoring the total number of photons at the deep Raman layer across all seven concentrations reveals a highly linear relationship, achieving an R^2^ value of 0.99, as shown in Fig. 10(b). To evaluate the model’s stability under varying acquisition times, spectra from a 100 mM guanine sample were acquired using integration times ranging from 10 to 60 s. Figure 10(c) compares the Raman signal recovered by our network with the baseline-corrected spectrum. The recovered signal closely tracks the baseline-corrected data, successfully resolving the same major peaks identified during the concentration analysis. Furthermore, the photon counts extracted from the deep Raman layer across the six acquisition intervals demonstrate a robust linear trend, yielding an R^2^ value of ∼0.99. Crucially, this demonstrates the network’s capacity to operate effectively under significantly shorter acquisition times, directly accelerating the overall high-throughput screening capabilities of the RamanBot platform. ^7^

**Figure 10:**
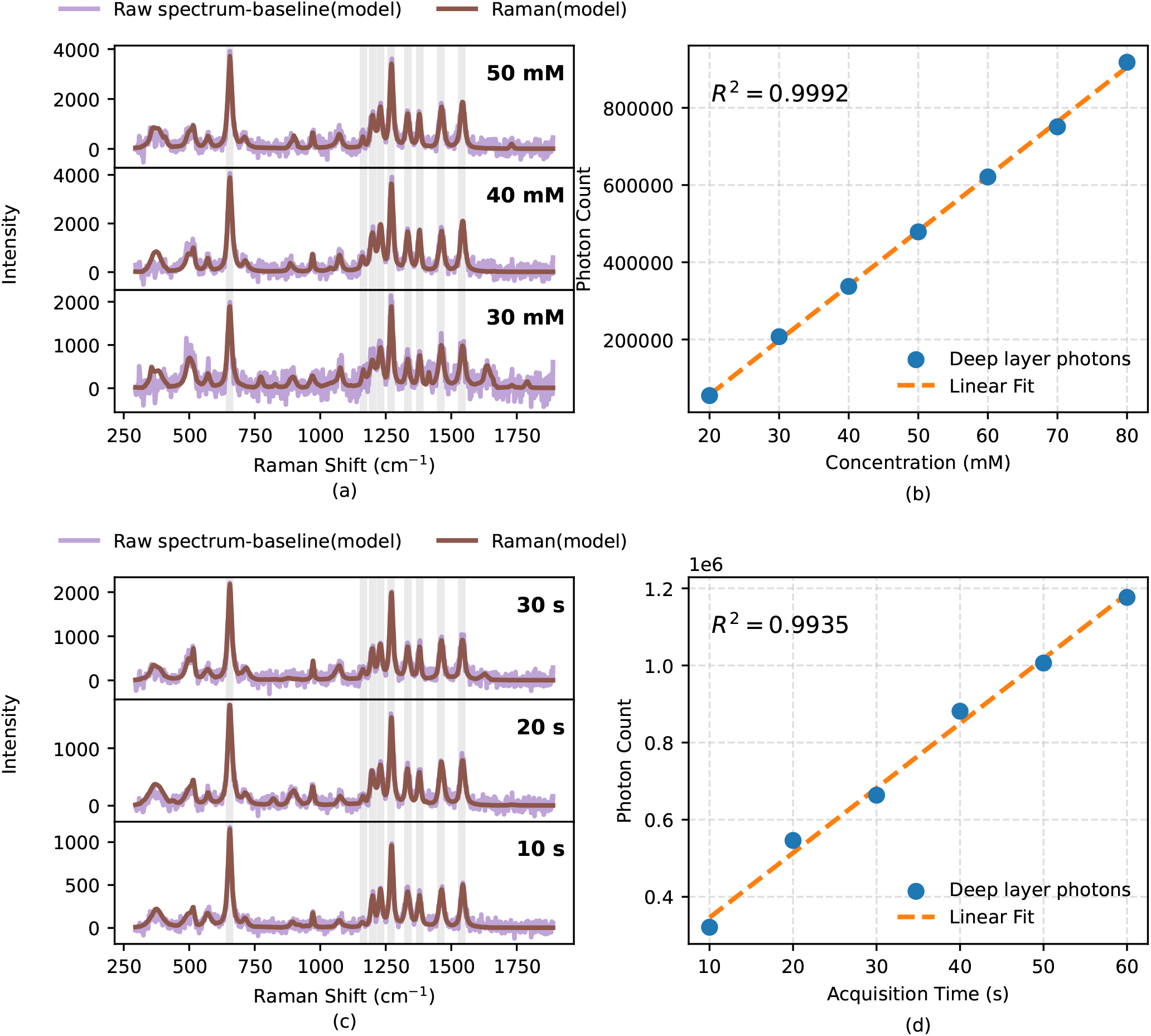
Denoising performance on guanine samples. (a) The recovered Raman signal in comparison with baseline corrected spectrum using our model for 7 different concentrations from 20 to 80 mM. The shaded columns represent the reported peaks at 654, 1165, 1200, 1230, 1270, 1335, 1380, 1460, and 1540 cm^−1^. (b) The integration of the photons at deep Raman layer monitored at different concentrations (c) The recovered Raman signal in comparison with baseline corrected spectrum using our model for 6 different acquisition time from 10 to 60s. (d) The integration of the photons at deep Raman layer monitored at different acquisition times.

## Conclusion

In conclusion, we have introduced a novel dual-branch U-Net architecture designed for the simultaneous baseline correction and denoising of Raman spectra. By utilizing a shared encoder that diverges at the bottleneck into dedicated baseline and Raman recovery decoding heads, our model effectively avoids distortions arise from sequentially applying denoising and baseline correction techniques. A key innovation of this architecture is the integration of a cross-attention gating mechanism, which provides spatial peak localization hints by evaluating feature differences between the encoder and the baseline decoder.

The model was trained using a custom-developed synthetic Raman engine configured to simulate spectra acquired by the RamanBot microplate platform. Comprehensive validation on these synthetic datasets demonstrated robust peak recovery across a broad range of signal-to-noise ratios (SNRs), preserving high spectral fidelity even under severe noise conditions.

We have successfully validated the model on experimental data. The network exhibited exceptional signal retrieval on highly noisy glycerol samples, and on moderately noisy adenine sulfate samples. Furthermore, our results confirmed that the network preserves relative peak intensities, enabling reliable quantitative analysis. When benchmarked against conventional baseline correction algorithms and spectral smoothing filters, our dual-head approach consistently demonstrated superior performance. Finally, counting deep Raman layer photons has shown perfect R^2^ value of 0.99 for both quantitative analysis and across different acquisition times. These results demonstrate that our model can significantly improve the SNR values and reduce the acquisition time leading to faster screening speed.

## References

(1) Buckley, K.; Matousek, P. Raman spectroscopy in pharmaceutical analysis: A review. Applied Spectroscopy 2017, 71, 2235–2252.

(2) Ferrari, A. C.; Basko, D. M. Raman spectroscopy as a versatile tool for studying the properties of graphene. Nature Nanotechnology 2013, 8, 235–246.

(3) Ebenezer, J.; et al. Raman spectroscopy: A potential tool for medical diagnostics and monitoring. Journal of Biophotonics 2021, 14, e202100103.

(4) Xu, G.; et al. Raman spectroscopy for the identification of microplastics in the marine environment. Marine Pollution Bulletin 2019, 146, 584–597.

(5) Kardash, E.; others Raman spectroscopy: A versatile tool in modern analytical chemistry. Analytical Chemistry 2023, 95, 120–135.

(6) Goldrick, S.; Umprecht, A.; Tang, A.; Zakrzewski, R.; Cheeks, M.; Turner, R.; Charles, A.; Les, K.; Hulley, M.; Spencer, C.; others High-throughput Raman spectroscopy combined with innovate data analysis workflow to enhance bio-pharmaceutical process development. Processes 2020, 8, 1179.

(7) Atia, K.; Hunter, R.; Asare-Werehene, M.; Tsang, B. K.; Anis, H. RamanBot: Versatile high throughput Raman system. PLoS One 2026, 21, e0334679.

(8) McCreery, R. L. Raman Spectroscopy for Chemical Analysis; Chemical Analysis: A Series of Monographs on Analytical Chemistry and Its Applications; John Wiley & Sons, Inc., 2000; Vol. 157.

(9) Wei, D.; Chen, S.; Liu, Q. Raman spectroscopy of biological tissues: Background mitigation and signal-to-noise optimization. Journal of Biomedical Optics 2015, 20, 111210.

(10) Savitzky, A.; Golay, M. J. Smoothing and differentiation of data by simplified least squares procedures. Analytical Chemistry 1964, 36, 1627–1639.

(11) Zhao, J.; Lui, H.; McLean, D. I.; Zeng, H. Wavelet transform-based denoising of Raman spectra: A comprehensive review. Journal of Raman Spectroscopy 2021, 52, 555–568.

(12) Zhang, Z.-M.; Chen, S.; Liang, Y.-Z. Baseline correction for Raman spectra using an adaptive iteratively reweighted penalized least squares strategy. The Analyst 2010, 135, 1138–1146.

(13) Brandt, J.; Mattsson, K.; Hassellöv, M. Deep Learning for Reconstructing Low-Quality FTIR and Raman Spectra—A Case Study in Microplastic Analyses. Analytical Chemistry 2021, 93, 16360–16368.

(14) Kazemzadeh, M.; Martinez-Calderon, M.; Xu, W.; Chamley, L. W.; Hisey, C. L.; Broderick, N. G. Cascaded deep convolutional neural networks as improved methods of preprocessing raman spectroscopy data. Analytical chemistry 2022, 94, 12907–12918.

(15) Fan, X.; others Deep learning-based Raman spectroscopy denoising and feature extraction. Analyst 2020, 145, 3551–3559.

(16) Luo, R.; Popp, J.; Bocklitz, T. Deep learning for Raman spectroscopy: A review. Analytica 2022, 3, 287–301.

(17) Wu, S.; Zhang, Y.; He, C.; Luo, Z.; Chen, Z.; Ye, J. Self-Supervised Learning for Generic Raman Spectrum Denoising. Analytical Chemistry 2024, 96, 17476–17485.

(18) Loc, I.; Kecoglu, I.; Unlu, M. B.; Parlatan, U. Denoising Raman spectra using fully convolutional encoder–decoder network. Journal of Raman Spectroscopy 2022, 53, 1445–1452.

(19) Ma, X.; Wang, K.; Chou, K. C.; Li, Q.; Lu, X. Conditional generative adversarial network for spectral recovery to accelerate single-cell Raman spectroscopic analysis. Analytical Chemistry 2022, 94, 577–582.

(20) Huang, J.; Zhou, F.; Cai, C.; Chu, R.; Zhang, Z.; Liu, Y. Remote SERS detection at a 10-m scale using silica fiber SERS probes coupled with a convolutional neural network. Optics Letters 2023, 48, 896–899.

(21) Sjöberg, J.; Siminea, N.; Păun, A.; Lita, A.; Larion, M.; Petre, I. Radar: raman spectral analysis using deep learning for artifact removal. Advanced Optical Materials 2025, 13, 2500736.

(22) Gao, C.; Zhao, P.; Fan, Q.; Jing, H.; Dang, R.; Sun, W.; Feng, Y.; Hu, B.; Wang, Q. Deep neural network: As the novel pipelines in multiple preprocessing for Raman spectroscopy. Spectrochimica Acta Part A: Molecular and Biomolecular Spectroscopy 2023, 302, 123086.

(23) Sperber, M.; Neubig, G.; Niehues, J.; Waibel, A. Attention-passing models for robust and data-efficient end-to-end speech translation. Transactions of the Association for Computational Linguistics 2019, 7, 313–325.

(24) Wang, Z.; D’Amico, A.; Borraccini, G.; Raj, A.; Huang, Y.-K.; Han, S.; Wang, T.; Ruffini, M.; Kilper, D.; Chen, T. Scalable ML models and cascaded learning for efficient multi-span OSNR and GSNR prediction. Journal of Optical Communications and Networking 2025, 18, A88–A99.

(25) Torbarina, L.; Ferkovic, T.; Roguski, L.; Mihelcic, V.; Sarlija, B.; Kraljevic, Z. Challenges and opportunities of using transformer-based multi-task learning in NLP through ML lifecycle: A position paper. Natural Language Processing Journal 2024, 7, 100076.

(26) Ruder, S. An overview of multi-task learning in deep neural networks. arXiv preprint arXiv:1706.05098 2017,

(27) Suteu, M.; Guo, Y. Regularizing deep multi-task networks using orthogonal gradients. arXiv preprint arXiv:1912.06844 2019,

(28) Chen, R. T.; Rubanova, Y.; Bettencourt, J.; Duvenaud, D. K. Neural ordinary differential equations. Advances in neural information processing systems 2018, 31 .

(29) Huang, S.; Hu, L.; Cheng, Z.; Su, S.; Wei, J.; Tong, X.; Zhao, Z. UDCNet: A U-Net Guided Dual-Branch Cross-Attention Network for SAR Object Detection. IEEE Journal of Selected Topics in Applied Earth Observations and Remote Sensing 2025,

(30) Lee, C.-Y.; Xie, S.; Gallagher, P.; Zhang, Z.; Tu, Z. Deeply-supervised nets. Artificial intelligence and statistics. 2015; pp 562–570.

(31) Szegedy, C.; Liu, W.; Jia, Y.; Sermanet, P.; Reed, S.; Anguelov, D.; Erhan, D.; Vanhoucke, V.; Rabinovich, A. Going deeper with convolutions. Proceedings of the IEEE conference on computer vision and pattern recognition. 2015; pp 1–9.

(32) Atia, K.; Hunter, R.; Asare-Werehene, M.; Tsang, B. K.; Anis, H. A programmable Raman plate reader based on a modified 3D printer for high-throughput screening. Optical Diagnostics and Sensing XXVI: Toward Point-of-Care Diagnostics. 2026; pp 57–63.

(33) Andor Technology HoloSpec On-axis high throughput imaging spectrograph. Oxford Instruments, 2025.

(34) Massie, C.; Chen, K.; Berger, A. J. Calibration technique for suppressing residual etalon artifacts in slit-averaged Raman spectroscopy. Applied Spectroscopy 2022, 76, 115–120.

(35) Zhao, J.; Lui, H.; McLean, D. I.; Zeng, H. Automated autofluorescence background subtraction algorithm for biomedical Raman spectroscopy. Applied spectroscopy 2007, 61, 1225–1232.

(36) Kim, B.; Chudomelka, B.; Park, J.; Kang, J.; Hong, Y.; Kim, H. J. Robust neural networks inspired by strong stability preserving Runge-Kutta methods. European Conference on Computer Vision. 2020; pp 416–432.

(37) Zhu, M.; Chang, B.; Fu, C. Convolutional neural networks combined with Runge– Kutta methods. Neural Computing and Applications 2023, 35, 1629–1643.

(38) Bousmalis, K.; Trigeorgis, G.; Silberman, N.; Krishnan, D.; Erhan, D. Domain separation networks. Advances in neural information processing systems 2016, 29.

(39) Young, C.; Liu, J.; Mortensen, M. L.; Feng, Y.; Li, E.; Wang, Z.; Guo, X.; Rosso, K. M.; Zhang, X. OASIS: A Deep Learning Framework for Universal Spectroscopic Analysis Driven by Novel Loss Functions. 2025; https://arxiv.org/abs/2509.11499.

(40) Diederik, K. Adam: A method for stochastic optimization. (No Title) 2014,

(41) Gryniewicz-Ruzicka, C. M.; Arzhantsev, S.; Pelster, L. N.; Kauffman, J. F.; Lucarelli, C.; Buhse, L. F. Multivariate calibration and instrument standardization for the rapid detection of diethylene glycol in glycerin by Raman spectroscopy. Applied Spectroscopy 2011, 65, 334–341.

(42) Mudalige, A.; Pemberton, J. E. Raman spectroscopy of glycerol/D2O solutions. Vibrational Spectroscopy 2007, 45, 27–35.

(43) Thurn, R.; Kiefer, W. Structural resonances observed in the Raman spectra of optically levitated liquid droplets. Applied Optics 1985, 24, 1515–1519.

(44) Lord, R. C.; Thomas, G. J. Raman spectral studies of nucleic acids and related molecules—I Ribonucleic acid derivatives. Spectrochimica Acta Part A: Molecular Spectroscopy 1967, 23, 2551–2591.

(45) Periasamy, A. Vibrational studies of Na 2 SO 4, K 2 SO 4, NaHSO 4 and KHSO 4 crystals. 2009,

(46) Madzharova, F.; Heiner, Z.; Kneipp, J. Surface-Enhanced Hyper-Raman Spectra of Adenine, Guanine, Cytosine, Thymine, and Uracil. The Journal of Physical Chemistry C 2017, 121, 1235–1242.

(47) Madzharova, F.; Heiner, Z.; Guhlke, M.; Kneipp, J. Surface-enhanced hyper-Raman spectra of adenine, guanine, cytosine, thymine, and uracil. The Journal of Physical Chemistry C 2016, 120, 15415–15423.

(48) Safar, W.; Azziz, A.; Edely, M.; Lamy de la Chapelle, M. Conventional Raman, SERS and TERS studies of DNA compounds. Chemosensors 2023, 11, 399.

